# Behavioral effect of chemogenetic inhibition is directly related to receptor transduction levels in rhesus monkeys

**DOI:** 10.1101/331694

**Authors:** Nicholas A. Upright, Stephen W. Brookshire, Wendy Schnebelen, Christienne G. Damatac, Patrick R. Hof, Philip G. F. Browning, Paula L. Croxson, Peter H. Rudebeck, Mark G. Baxter

**Affiliations:** Fishberg Department of Neuroscience and Friedman Brain Institute, Icahn School of Medicine at Mount Sinai, New York, New York, 10029

**Author notes:** Contributed equally to this study (Nicholas A. Upright, Stephen W. Brookshire, Wendy Schnebelen).

## Abstract

We used inhibitory DREADDs (Designer Receptors Exclusively Activated by Designer Drugs) to reversibly disrupt dorsolateral prefrontal cortex (dlPFC) function in male macaque monkeys. Monkeys were tested on a spatial delayed response task to assess working memory function after intramuscular injection of either clozapine-*N*-oxide (CNO) or vehicle. CNO injections given before DREADD transduction were without effect on behavior. rAAV5/hsyn-hM4Di-mCherry was injected bilaterally into the dlPFC of five male rhesus monkeys, to produce neuronal expression of the inhibitory (Gi-coupled) DREADD receptor. We quantified the percentage of DREADD-transduced cells using stereological analysis of mCherry-immunolabeled cells. We found a greater number of immunolabeled neurons in monkeys that displayed CNO-induced behavioral impairment after DREADD transduction compared to monkeys that showed no behavioral effect after CNO. Even in monkeys that showed reliable effects of CNO on behavior after DREADD transduction, the number of prefrontal neurons transduced with DREADD receptor was on the order of 3% of total prefrontal neurons counted. This level of histological analysis facilitates our understanding of behavioral effects, or lack thereof, after DREADD vector injection in monkeys. It also implies that a functional silencing of a relatively small fraction of dlPFC neurons, albeit in a widely distributed area, is sufficient to disrupt spatial working memory.

**Significance Statement:** Cognitive domains such as working memory and executive function are mediated by the dorsolateral prefrontal cortex (dlPFC). Impairments in these domains are common in neurodegenerative diseases as well as normal aging. The present study sought to measure deficits in a spatial delayed response task following activation of viral-vector transduced inhibitory DREADD (Designer Receptor Exclusively Activated by Designer Drug) receptors in rhesus macaques and compare this to the level of transduction in dlPFC using stereology. We found a significant relationship between the extent of DREADD transduction and the magnitude of behavioral deficit following administration of the DREADD actuator compound clozapine-*N*- oxide (CNO). These results demonstrate it will be critical to validate transduction to ensure DREADDs remain a powerful tool for neuronal disruption.

## Introduction

Chemogenetic techniques such as DREADDs (Designer Receptors Exclusively Activated by Designer Drugs) allow for the remote manipulation of neuronal activity. They can be targeted to distinct cell populations defined by anatomical, connectional, or other phenotypic characteristics and are activated by systemic administration of an otherwise inert drug (Armbruster et al., 2007); (Roth, 2016). The most commonly used DREADD system employs the clozapine metabolite clozapine-N-oxide (CNO) to activate a modified muscarinic acetylcholine receptor which is no longer sensitive to acetylcholine as an agonist (Armbruster et al., 2007). In principle, CNO has no endogenous receptors or other physiological effects and is inert in the absence of the DREADD receptor, and because the DREADD receptor has no endogenous ligand it is inert until CNO or another DREADD actuator compound is administered. The DREADD receptor can be linked to different G-protein signal transduction mechanisms, meaning that it is possible to stimulate or inhibit neuronal activity via this system (Roth, 2016).

Chemogenetic techniques would be particularly powerful if implemented in nonhuman primates. They would allow for experimental designs with reversible manipulation of neuronal activity across large anatomical regions that are beyond the reach of electrical or optogenetic stimulation (Ohayon et al., 2013) and on time scales consistent with cognitive testing or behavioral neurophysiology. The success of these techniques in modifying behavior when DREADD receptors are expressed in rostromedial caudate (Nagai et al., 2016) or orbitofrontal cortex (Eldridge et al., 2015) has been demonstrated. To effectively employ these technologies, there is a need for studies that not only demonstrate the efficacy of DREADDs in non-human primates, but also validate the technique and illustrate challenges to address in the future. Our goals in the present study were twofold. First, we sought to implement DREADDs in nonhuman primates in a different neocortical area, the dorsolateral prefrontal cortex (dlPFC), using a sensitive behavioral probe of dlPFC function, the spatial delayed response task (Goldman and Rosvold, 1970; Bachevalier and Mishkin, 1986). Second, upon observing variability in outcomes after DREADD-bearing viral vector injections into dlPFC, we applied unbiased stereological counting techniques to postmortem histological analysis of dlPFC in order to determine the relationship between the extent of DREADD receptor transduction within dlPFC with the magnitude of behavioral effect caused by DREADD receptor activation. As these studies extended over a period of several years and additional information became available about the use of chemogenetic techniques in monkeys, we incorporated additional control conditions into our design.

## Materials and Methods

### Subjects

Five male rhesus macaques, denoted cases A, B, P, T, and Z, aged between 44 and 76 months old and weighing 4.5 to 9.4 kg at the time of surgery, were used for this study. Monkeys were socially housed indoors in single sex groups. Daily meals, consisting of a ration of monkey chow and a variety of fruits and vegetables, was given within transport cages once testing was completed, except on weekends when they were fed in their home cages. Within the home cages water was available ad libitum. Environmental enrichment, in the form of play objects or small food items, was provided daily in the home cages. All procedures were approved by the Icahn School of Medicine Institutional Animal Care and Use Committee and conform to NIH guidelines on the use of non-human primates in research.

### Apparatus

Testing was performed within a Wisconsin General Testing Apparatus (WGTA). The WGTA is a small enclosed testing area where the experimenter can manually interact with the monkey during testing. Monkeys were trained to move from the home cage enclosure to a metal transport cage, which was wheeled into the WGTA. The experimenter was hidden from the monkey’s view by a one-way mirror, with only the experimenter’s hands visible. A sliding tray with two food wells could be advanced within reach of the monkey, with a pulley-operated opaque black screen that could be lowered to separate the tray from the monkey.

### Behavioral Testing

Training on the delayed response task followed Bachevalier and Mishkin (1986) and Croxson et al. (2011). Monkeys were shown a small food reward (a peanut, M&M, raisin, or craisin, depending on each monkey’s preference), which was placed in one of two food wells on a test tray. The left/right location of the reward across trials was always determined based on a pseudorandomized, counterbalanced sequence. An opaque black screen was then lowered between test tray and the monkey for a predefined delay period. The screen was subsequently raised and the test tray was advanced within reach of the monkey. During initial shaping, the monkeys were taught to displace a flat, gray tile covering one of the two small food wells. Once monkeys readily displaced the tiles, they were advanced along three stages of training, with 24- 30 trials per session. Once each monkey successfully completed the third stage of training to criterion, experimental testing began, consisting of sessions of 24 trials. On each trial, if the monkey reached for the correct well it was allowed to take the reward, otherwise the tray was quickly pulled back before the monkey could reach for the other well.

For the first stage of training, both the baited and non-baited wells were covered, and the tray advanced for the monkey’s choice. Trials were repeated until the monkey chose correctly. The second stage included a brief (“0 second”) lowering of the opaque black screen between the monkey and the food tray (the screen was lowered and then raised immediately) before the tray was presented to the monkey for choice. Each monkey advanced from the first to the second stage, and from the second to the third, after completing two consecutive sessions at 90% correct or better. The third stage was the same as the second with longer delays (1-5 s) of the opaque screen. Each monkey started at a 1-s delay and was advanced to a 1-s longer delay upon 90% correct or better performance, and was reduced 1-s (to a minimum of 1-s) upon performance of less than 90% correct. Once 90% correct performance or better was achieved in one session at the 5-s delay, training was complete and experimental testing began.

In the experimental task, each trial consisted of one of four possible delays (5, 10, 15, and 20 s) varied pseudorandomly across trials such that each delay occurred 6 times in each test session. For case T, 5 delays were used (5, 10, 15, 20, 30 s), each occurring 6 times in each test session. Monkeys continued with the variable delay task for the remainder of the experiment. Drug injections began once each monkey reached stable performance on the experimental task. Typically, drug injections were given one or two days per week, with vehicle or no-injection testing on other days, and at least one day of rest given after CNO injections. Training on the delayed response task was conducted after surgery in two cases (A and B) and before in the three others (cases P, T, and Z).

### Drugs

Clozapine-N-oxide (CNO) was obtained through the National Institute of Mental Health Chemical Synthesis Program. DMSO was obtained from Sigma, and 0.1 M phosphate buffered saline (PBS) was prepared in-house. CNO was first dissolved in DMSO, and then diluted in PBS to a final concentration of 15 mg/mL in 15/85 DMSO/PBS (v/v). Alternately, CNO was converted to the hydrochloride salt CNO HCl in the laboratory of Dr. Jian Jin (Icahn School of Medicine at Mount Sinai). Compound 21 was obtained from Dr. Jian Jin. For administration, these drugs were weighed to compose the appropriate dose for each monkey and dissolved in 1.5 ml PBS. The final solutions were filtered through a 0.2 µm syringe filter before injection. CNO/CNO HCl was given at 10 or 20 mg/kg, i.m. During drug testing, monkeys received 240 mg Vitamin C orally each day (chewable supplements), in an effort to inhibit hepatic conversion of CNO to clozapine (Pirmohamed et al., 1995). Drug injections were administered one hour prior to the behavioral session on testing days. Because of the large volume of CNO injections required to achieve 20 mg/kg dose, injections were split between several i.m. injection sites. To maintain behavioral performance while limiting the number of times monkeys had to receive injections, vehicle injection sessions were interspersed with test sessions that were not preceded by injection. The number of CNO injection sessions for each monkey was limited mainly by availability of CNO, owing to the large doses required for each CNO test session. **Table 1** shows the schedule of injections and CNO dosing that each animal received during the course of post-operative testing.

**Table 1.** Surgery and injection schedule during the course of post-operative behavioral testing

| Case | Surgery Date | CNO (20 mg/kg) Test Session |  |  |  |
| --- | --- | --- | --- | --- | --- |
|  |  | 1 | 2 | 3 | 4 |
| A | 12/11/13 | 11/04/14 | 11/25/14 |  |  |
| B | 12/10/13 | 11/04/14 | 11/07/14 | 11/25/14 | 12/10/14 |
| P | 04/09/15 | 05/19/15 | 05/21/15 | 07/21/15 |  |
| T | 08/02/16 | 10/06/16 | 11/06/16 | 12/13/16 |  |
| Z | 04/07/15 | 05/19/15 | 05/21/15 |  |  |

### Surgery

Surgical methods for bilateral injection of DREADD AAV into the dlPFC followed Croxson et al. (2011). Aliquots (100 µl) of rAAV5/hsyn-hM4Di-mCherry were obtained from the University of North Carolina at Chapel Hill Vector Core at 2.4 (cases A, B), 3.5 (case T), or 4.1 ×; 10^12^ (cases P, Z) particles/ml. AAV was stored at −80° C until immediately before injection, at which time it was thawed on wet ice and loaded into 10 µl Hamilton syringes for intracortical injection. A new aliquot of AAV was used for each surgical procedure (1 or 2, depending on the number of injections performed) and leftover AAV was not refrozen. Surgical procedures took place in a dedicated operating suite under strict aseptic conditions. On the day of surgery, each monkey was sedated with ketamine (10 mg/kg) and dexmedetomidine (0.01 mg/kg), transported to the surgical preparation area, the head shaved and cleansed, and intubated with an endotracheal tube. Anesthesia was maintained with sevoflurane (2-4%, to effect) in 100% oxygen. The skin and galea were incised and retracted, a single bone flap turned over the frontal lobes bilaterally, and the dura reflected over the dlPFC in each hemisphere. Injections (1 µl) of AAV were made into each hemisphere under visual guidance through an operating microscope, with care taken to place the beveled tip of the Hamilton syringe containing the AAV at an oblique angle to the pial surface. Injections were spaced approximately 2 mm apart and covered the dorsal and ventral borders of the principal sulcus, extending dorsally towards the midline bounded anteriorly by the tip of the principal sulcus, posteriorly by a line connecting the posterior tip of the principal sulcus and the anterior tip of the ascending limb of the arcuate sulcus, and dorsally by an approximate line extending from the anterior tip of the arcuate sulcus anteriorly towards the front of the skull, parallel to the midline. Each case received injections in the left and right hemispheres as follows: Case A, 49 and 54; Case B, 57 and 50; Case P, 45 and 66; Case T, 36 and 53; Case Z, 86 and 98. Variation in number of injections was related to individual differences in brain size, sulcal morphology, and degree of exposure. Upon completion of injections in each hemisphere the dura was closed with Vicryl sutures; when injections in both hemispheres were completed the bone flap was replaced and held in place with loose Vicryl sutures, the galea and skin were closed in layers, and anesthesia was discontinued. The monkey was extubated when a swallowing reflex was present, returned to the home cage, and monitored continuously until normal motor behavior resumed. Each monkey remained individually housed for a few days after surgery until the attending veterinarian determined it could rejoin the social group. Postoperative treatment included buprenorphine (0.01 mg/kg i.m. every 8 hours) and meloxicam (0.2 mg/kg i.m. every 24 hours) for analgesia for 1-5 days and 3- 5 days respectively, based on veterinary guidance, as well as cefazolin (25 mg/kg i.m. every 24 hours) for 5 days and dexamethasone sodium phosphate (0.4-1 mg/kg, every 12-24 hours) for 5 days on a descending dose schedule. Postoperative behavioral data collection began a minimum of 2 weeks after surgery. CNO test sessions were carried out 324-349 days postoperatively in case A, 325-365 days in case B, 40-103 days in case P, 65-133 days in case T, and 42-44 days in case Z.

### Histology

#### Tissue Extraction

At the conclusion of behavioral testing, each monkey was sedated with ketamine (10 mg/kg) i.m., intubated, and an intravenous line placed for i.v. administration of a terminal anesthetic dose of sodium pentobarbital (100 mg/kg). Upon loss of corneal reflex, the chest cavity was opened, the descending aorta clamped, and transcardial perfusion was initiated with 1% paraformaldehyde (PFA) for 90 seconds, and then 4% PFA for ∼15 minutes, via a cannula placed in the ascending aorta. Brains were extracted and placed in 4%PFA and kept at 4°C overnight, before being placed in sucrose/sodium azide solutions. The brains were cryoprotected in increasing concentrations of sucrose, 10%, 20% and 30%, with 0.1% of sodium azide added to all as a preservative agent. Brains were kept at 4°C in 30%sucrose/0.1% sodium azide until ready to be sectioned.

#### Sectioning

Brains were removed from the cryoprotectant solution and the brainstem and posterior part of the occipital lobe were blocked to given a level surface to sit the brain upright. A sliding microtome and freezing stage were used to section the brain. Coronal sections were cut at 50 µm thickness, and a 1:10 set was collected throughout the frontal lobes. Sets were collected into 0.1 M PBS w/ 0.1%azide for storage at 4°C, or into a cryoprotectant solution consisting of glycerol, ethylene glycol, PBS, and distilled water (30/30/10/30 v/v/v/v, respectively) for storage at −80°C.

#### Immunohistochemistry

Sections were taken from the 4°C storage sets and probed for mCherry immunoreactivity. Endogenous peroxidase activity was quenched with 0.3%hydrogen peroxide (v/v) with 20%methanol (v/v) in PBS. Alternating sections were blocked with 5%normal goat serum (VectaShield, S-1000), and then incubated in anti-mCherry rabbit polyclonal primary antibody in blocking solution (Abcam, ab167453; 1:50,000 dilution). The secondary consisted of a goat-anti-rabbit biotin-conjugated antibody (Jackson Immunoresearch; #111-065-003; 1:500 dilution) in 2% normal goat serum, followed by the VectaShield ABC Peroxidase kit (VectaShield, PK-6100) with the modifications to the ABC kit instructions; 2 drops solution A and 2 drops solution B per 10 ml PBS with 0.1% Triton X-100. Finally, labeling was visualized using the DAB kit (VectaShield, SK-4100) with nickel enhancement according to manufacturer’s recommendations. Sections were counterstained with cresyl violet and mounted with DPX.

### Stereology

The dlPFC region of interest was identified using a 5x objective. The borders of the dlPFC were defined as the dorsal edge of the cingulate sulcus medially, the ventral lip of the principal sulcus laterally, and the anterior and posterior tips of the principal sulcus anteriorly and posteriorly. This region encompassed most of area 46 and 9/46d, and portions of areas 9/46v, 8b, 8Ad, and 9, as defined by Petrides and Pandya (1999). As such it approximated the dlPFC ablations made in other studies (Baxter et al., 2008; Bachevalier and Mishkin, 1986) and included more cortex than was targeted intraoperatively, because our injections did not extend into the midline.

Standard unbiased stereology was performed on sections using a Zeiss Apotome.2 light microscope equipped with a Q-Imaging digital camera, a motorized stage, and Stereo Investigator software (MBF Bioscience, Williston, VT). Cresyl violet and mCherry-positive cells were identified using the soma as the counting target and numbers were estimated using the optical fractionator probe following the process described in West et al. (1991). Pilot studies were performed to determine appropriately sized sampling grids and counting frames. Six to seven sections were used for each animal. Both cresyl and mCherry-stained tissue were counted using a x40 oil-immersion objective lens within a counting frame of 100 ×; 100 ×; 10 µm^3^ with the dissector top guard volume extended 1 µm below the tissue section surface. The sampling grid for cresyl cells was 700 ×; 700 µm^2^ and the sampling grid for mCherry-positive cells was 400 ×; 400 µm^2^. The cell body was used as the counting object for cells that fell within the dissector or across its inclusion planes. Neurons were differentiated from glia by cell morphology, identified with the cresyl violet stain, and the presence of a well-defined nucleolus. Coefficients of error were calculated for each region to ensure minimal variance due to sampling.

Representative histological images are shown in **Figure 1**. mCherry immunostaining was patchy, as expected. We did not quantify mCherry staining outside of the dlPFC; however, we observed some staining outside the region of interest in four out of the five cases. These stained cells may be attributed to unintended deeper penetrations during surgery as a result of the the handheld syringe technique. Cases A, B, and Z showed small patches of mCherry-positive cells in medial prefrontal cortex, mostly localized around the cingulate sulcus. Cases A and P showed several instances of stained cells in orbital prefrontal cortex, concentrated around the medial orbital sulcus region. No case exhibited staining in thalamus and posterior parietal cortex. Although four out of the five cases demonstrated variable degrees of ectopic staining, there was no obvious relationship between this ectopic staining and behavioral effects of CNO administration, with ectopic staining being present in the same cortical regions in both monkeys that showed effects of CNO administration post-surgery and those that did not.

**Figure 1.**
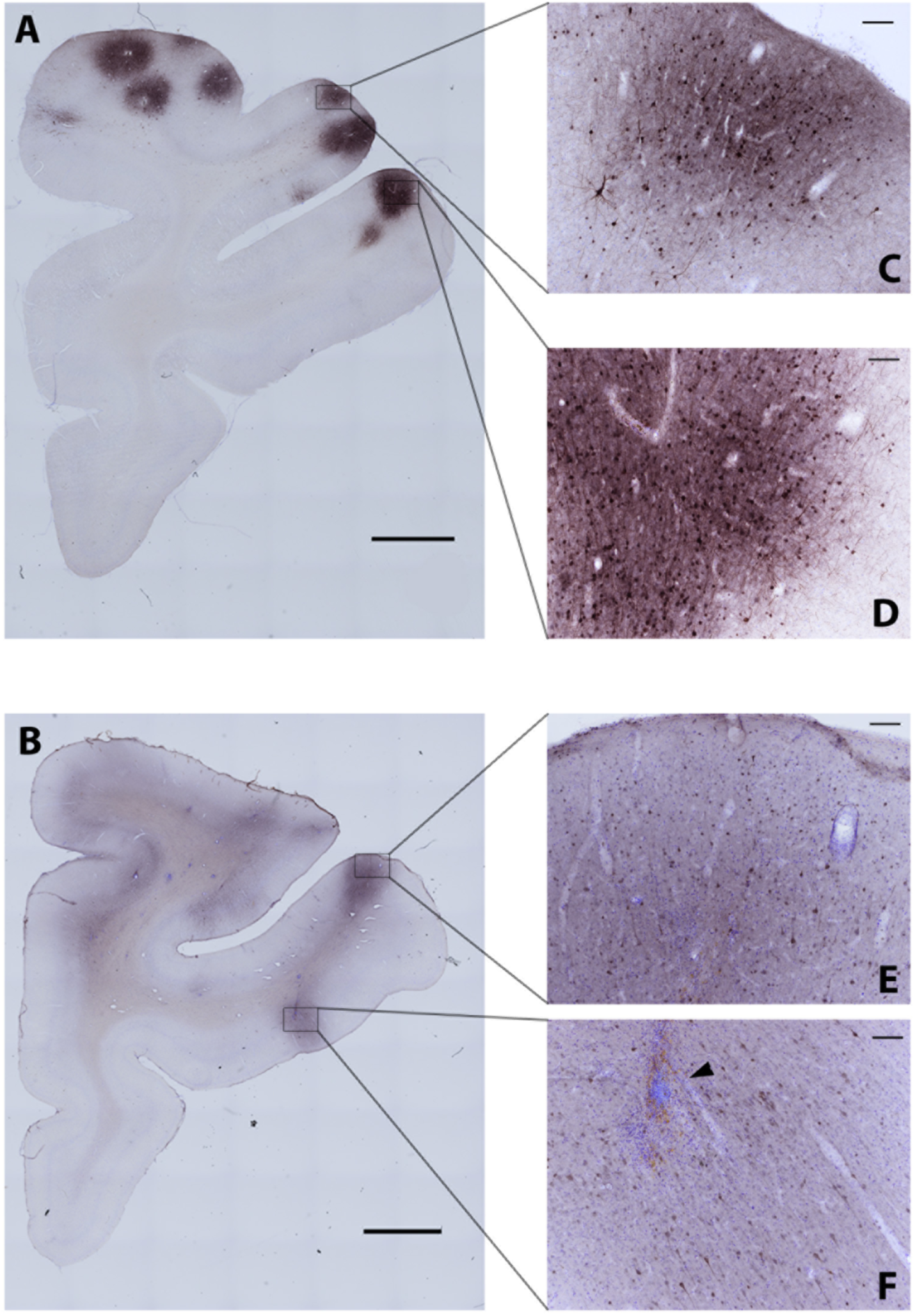
Transduction pattern in two example cases. ***A***, DREADD transduction pattern shown in PFC of Case P. Regions within black squares are shown at higher magnification in ***C*** and ***D***. ***B***, DREADD transduction from PFC of Case Z. ***C-D***, DREADD patches along principal sulcus from same slice as in ***A***. Shown at 10x magnification. ***E-F***, DREADD patches from same slice as in B. Shown at 10x magnification. Note DREADD-positive cells near principal sulcus (***E***) as well as orbital frontal cortex (***F***). Black arrowhead denotes presence of needle track. Scale bars: ***A*** and ***B***, 2500 µm; ***C-F***, 100 µm.

### Statistical Analysis

Because of variability across monkeys in the effect of CNO on performance after surgery, and because we were limited in the number of test sessions we could carry out with each monkey due to constraints on our supply of CNO and the large quantity of drug required to generate a sufficient dose for each behavioral test session, we adopted a case study-type approach to data analysis. For each monkey, we determined first the impact of vehicle injections and pre/post surgery on performance, using a 2 ×; 2 ANOVA on percent correct performance across vehicle and no-injection test sessions before and after surgery, treating each session as a unit of analysis within each monkey. No monkey showed an effect of vehicle injection relative to baseline sessions without vehicle injection, and for only one monkey (case Z) did performance differ between pre- and postoperative testing (performance was better after surgery than before). We used the single-case t-test approach of Crawford and Howell (1998) to evaluate each drug test session compared to postoperative vehicle injection sessions for each monkey, because the number and timing of drug injection test sessions varied across monkeys as we accumulated data and our experimental protocol evolved based on communication with other research groups that were carrying out studies with DREADDs in monkeys concurrently with ours. We evaluated significant impairments in drug testing sessions as one-tailed p-values < .05 generated from each t-test. Because we expected only impairments following drug administration a priori, we judged one-tailed tests to be appropriate. In any case, the evaluation of statistical significance of any individual test session is only a proxy for the absolute magnitude of the drug effect in terms of an increase in errors committed during the test session, which is readily apparent from the raw behavioral data.

To adjust for baseline differences in performance, we determined an aggregate “deficit score” based on postoperative DREADD receptor activation (Croxson et al., 2012). This score was calculated as a percent of maximal deficit in the spatial delayed response task, providing a single score characterizing each monkey’s behavioral impairment. We computed the correlation between this behavioral deficit score and the percentage of neurons in each monkey’s dlPFC that was transduced by DREADD receptors, based on the stereological quantification.

## Results

### The effects of DREADD-mediated dlPFC inhibition on spatial working memory

A summary of behavioral data for all five cases can be found in **Table 2**. Two monkeys demonstrated robust impairments in the spatial working memory task after injection of CNO (Cases B and P), one monkey demonstrated moderate impairment (Case T), and two monkeys showed no difference in spatial working memory function after CNO administration (Cases A and Z). It is interesting in Cases B and P that they were unimpaired during their final CNO test session, perhaps suggesting that with repeated dosing they were able to compensate for the impact of temporary neuronal inhibition following systemic CNO. Mean performance for vehicle and CNO sessions for each monkey is shown in **Figure 2**, both across all sessions in each condition (**Figure 2A**) and broken down by delay (**Figure 2B**). Given the limited number of total trials at each delay, we did not carry out data analyses at each delay individually, but it is interesting that case T appears to show deficits only at the longest delay tested.

**Table 2.**
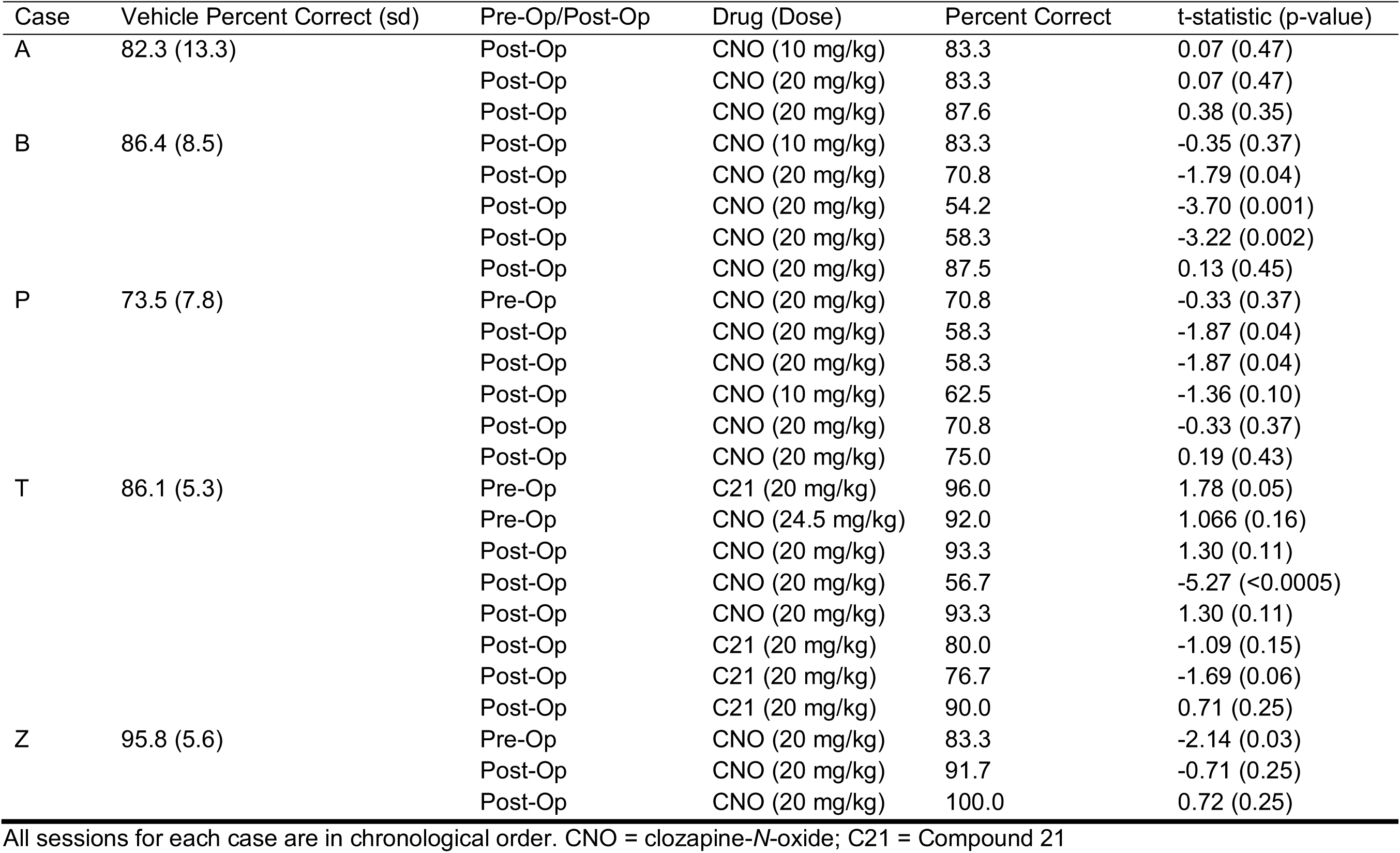
Behavioral data for pre-operative and post-operative testing sessions.

**Figure 2.**
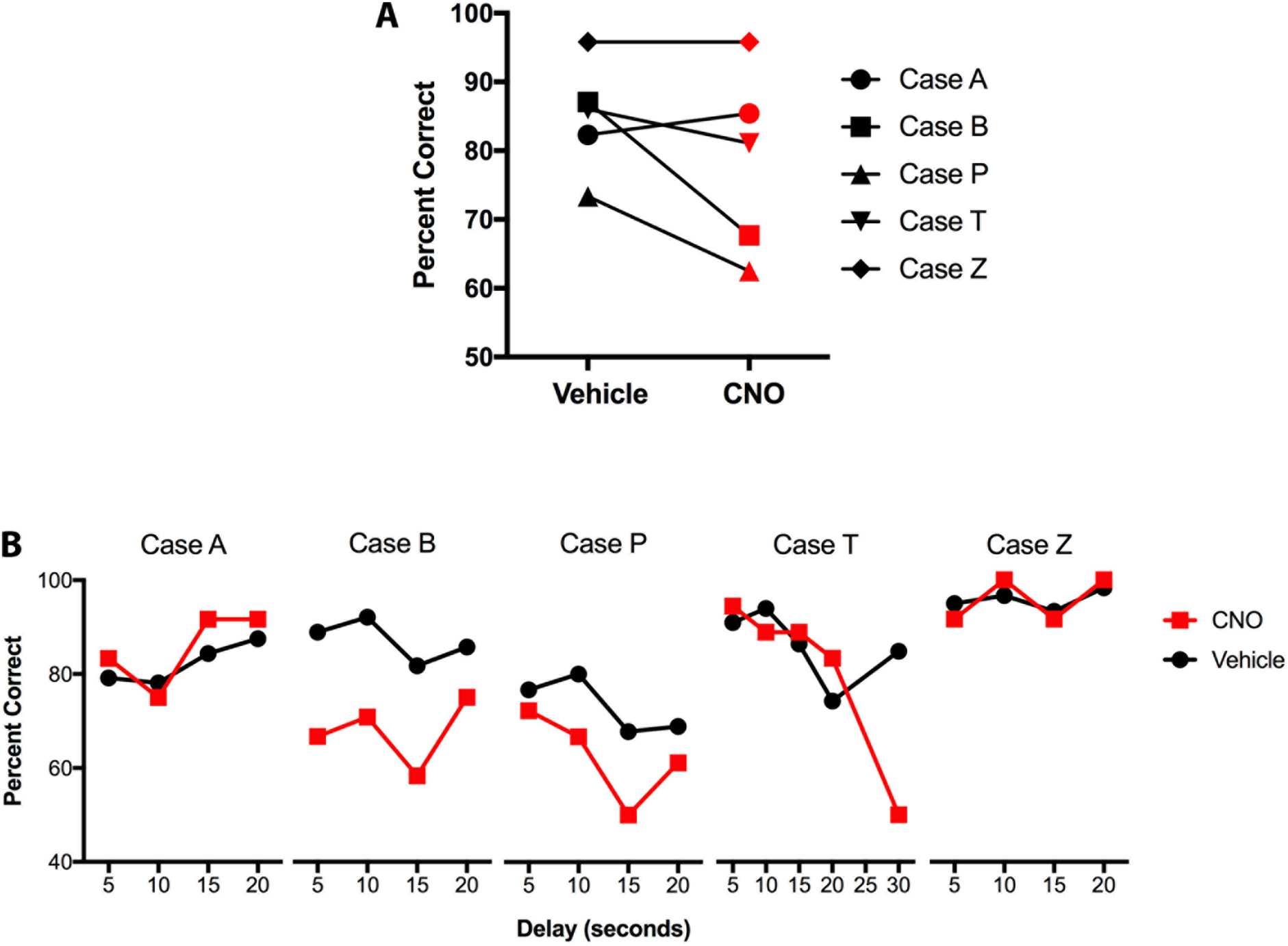
Spatial delayed response performance for vehicle and CNO sessions. ***A***, Mean performance across all sessions for each case. ***B***, Performance for each case broken down by delay (5, 10, 15, and 20 s). Case T had an additional trial time of 30 s.

### The effects of CNO on behavior

As a control measure for the effects of CNO on behavior, we obtained data on CNO injections prior to AAV injection surgery in two cases (one session each in cases P and Z). Case P, who showed marked effects of 20 mg/kg CNO after surgery, showed no effect of 20 mg/kg CNO preoperatively (70.8%correct 73.4% correct on vehicle, t = −0.33, p = 0.37. Case Z, who did not show effects of 20 mg/kg CNO after surgery, actually did worse on his pre-surgery CNO test compared to vehicle performance, 83.3% correct vs 95.8% correct on vehicle, t = −2.14, <u>p</u> =0.029, although this comparison is complicated by overall better postoperative performance compared to preoperative performance in this monkey, which elevates his mean vehicle score. In any case, this monkey showed no impairments in two further 20 mg/kg CNO sessions done after surgery, arguing against general nonspecific effects of CNO on behavior. Case T had one session with Compound 21 preoperatively, and scored 96% correct, suggesting Compound 21 also had no discernible behavioral effect in the absence of DREADD expression.

### The relationship between DREADD receptor transduction in dlPFC and spatial working memory performance

We conducted unbiased stereological counting on histological sections from the dlPFC of each animal to determine the relationship between behavioral performance in the spatial delayed response task and the proportion of neurons transduced with inhibitory DREADD receptor. Monkeys that demonstrated behavioral impairments showed an increased level of mCherry staining compared to monkeys that demonstrated no behavioral change after CNO injection. We found that there was a significant positive correlation between percent of positive mCherry-immunolabeled neurons in the dlPFC and performance on the spatial delayed response task after CNO administration calculated as a percent deficit score [(mean baseline - mean CNO)/(mean baseline - chance)], r = 0.8745, <u>p</u> = .0196, illustrated in **Figure 3.**

**Figure 3.**
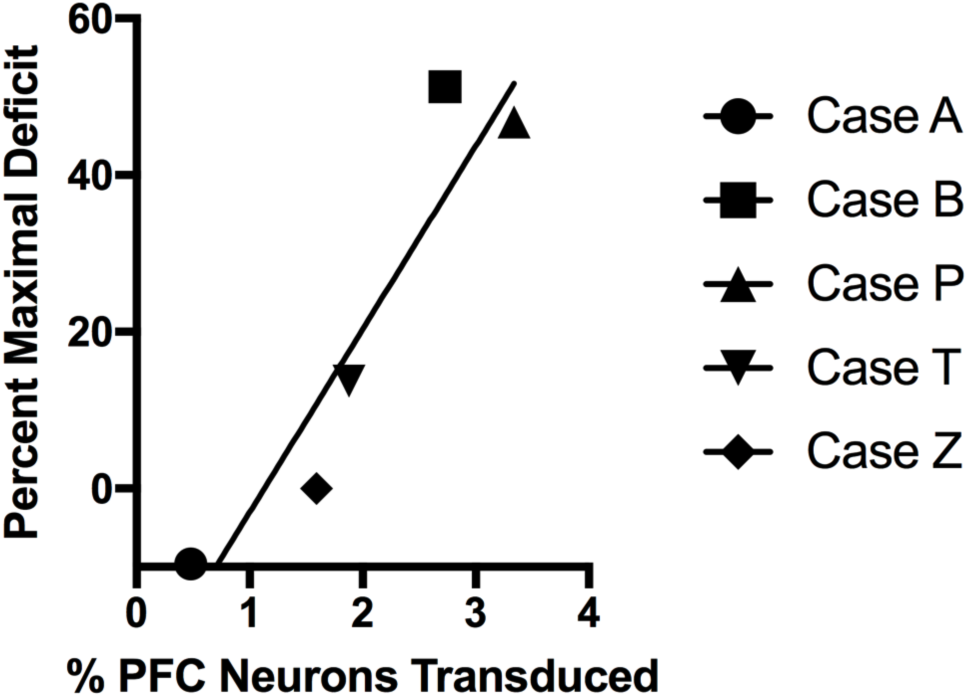
Significant positive correlation between stereological and behavioral measures. Percent of prefrontal neurons transduced with DREADD was significantly correlated with performance on the spatial delayed response task, calculated here as an aggregate deficit score from postoperative DREADD receptor activation (r = 0.8745, p = 0.0196).

## Discussion

In some monkeys who received hM4Di DREADD AAV injections into the dlPFC, we were able to obtain a reliable behavioral deficit in a spatial working memory task following systemic injections of CNO. In monkeys where we examined CNO injections preoperatively, no impact on behavior was seen. There was a monotonic relationship between the extent of DREADD receptor transduction within the dlPFC and the magnitude of behavioral impairment following DREADD receptor activation. Thus, there is a biological basis to apparently stochastic effects of DREADD transduction in rhesus monkeys. In the two monkeys with the largest behavioral deficits following DREADD receptor activation, only ∼3% of neurons within the dlPFC were transduced with DREADD receptors. These findings have implications for the implementation of chemogenetic approaches in nonhuman primates, as well as for the neurophysiology of cognitive functions of the prefrontal cortex.

Our goal in initiating this study was to determine, with a simple behavioral task dependent on a cortical area with a straightforward surgical approach, whether we could achieve a substantial behavioral deficit by activation of hM4Di DREADDs on par with what could be achieved by a cortical ablation or neurotoxic lesion. This would be a critical prelude to using chemogenetic methods in studies where the consequence of neuronal inhibition (or activation) would be less clearly determined a priori. We were only partially successful in accomplishing this goal. In two cases (B and P) we achieved substantial behavioral impairments with DREADD receptor activation, but in the other three cases, effects were equivocal. The basis of this variability was clearly related to the extent of DREADD receptor transduction within the dlPFC. Nagai et al. (2016) reported a similar phenomenon, where by in cases where systemic CNO failed to produce a behavioral deficit in their reward sensitivity task, they also did not observe significant displacement of ^11^C-clozapine in PET scans following CNO administration. Because they made this determination in vivo, they were able to make additional AAV injections in an effort to increase DREADD receptor expression and thereby achieve behavioral effects of CNO administration. Extent of DREADD receptor transduction appeared to be unrelated to surgical parameters such as number of injections in each case or lot of viral vector used. We did not determine the presence of AAV neutralizing antibodies in our monkeys prior to surgery; these have been reported after AAV injections into the central nervous system for delivery of optogenetic constructs (Mendoza et al., 2017), although it is not clear whether the presence of neutralizing antibodies would impede transduction by AAV (Gray et al., 2013).

Both monkeys that showed marked behavioral impairments after 20 mg/kg CNO appeared to adapt, with performance in each monkey’s final test session at this dose not differing significantly from baseline. Perhaps with a relatively small population of neurons in dlPFC being functionally silenced by systemic CNO, the monkeys were able to compensate after repeated behavioral testing on the delayed response task under CNO. This could not have been related to level of DREADD expression as a function of time post-transduction, because the interval between surgery and test for case B was quite long and was much briefer for case P. It has also been shown that significant desensitization of DREADDs apparently does not occur in vivo (Roth, 2016; Roman et al., 2016). Nonetheless, this may suggest some additional limitations on using functional silencing with chemogenetic techniques as a tool to investigate behavioral deficits after inhibitions of specific populations of neurons.

Although CNO is present in the cerebrospinal fluid (CSF) in monkeys (Eldridge et al., 2015; Raper et al., 2017), it has also become apparent recently that CNO is actively transported out of brain parenchyma by P-glycoprotein (Raper et al., 2017). Thus CSF concentrations may not reflect CNO availability at neuronally expressed DREADD receptors. Moreover, conversion of CNO to clozapine occurs in monkeys, producing concentrations of clozapine sufficient to bind to DREADD receptors (Raper et al., 2017). A recent study in rodents suggests that the mechanism of hM3/hM4 DREADD receptor activation is exclusively via conversion of CNO to clozapine (Gomez et al., 2017). We saw no effect of CNO injections on behavior in our monkeys prior to DREADD receptor expression so we do not think positive effects of CNO, where observed, are explained merely by clozapine interaction with non-DREADD receptors. It remains a logical possibility that individual differences in CNO penetration of the central nervous system, or of conversion of CNO to clozapine, influence the behavioral effectiveness of DREADD receptor activation. Because we did not determine CSF levels of CNO/clozapine in our monkeys, we cannot address this possibility. As suggested by Gomez et al. (2017), future studies might use low doses of clozapine, that do not produce behavioral effects on their own, as DREADD actuators rather than CNO. There is an alternative non-CNO actuator, compound 21 (Chen et al., 2015) that is a potential alternative to CNO/clozapine. We only had the opportunity to carry out limited experiments with compound 21 in this study, but it appears comparable to CNO in potency. Another possibility for future studies is using an alternative DREADD receptor system such as that based on modification of the kappa opioid receptor (KORD-DREADD; (Vardy et al., 2015). However, this approach has limited application in monkeys at the moment because the limited solubility of the KORD-DREADD ligand salvanorin B makes dosing impractical.

Despite the technical challenges encountered, we were able to impair spatial delayed response performance, a task dependent on intact dlPFC, by DREADD receptor activation in monkeys in which ∼3% of prefrontal neurons were transduced with the hM4Di DREADD receptor. This implies that inhibition of a relatively small fraction of dlPFC neurons is sufficient to produce substantial behavioral deficits in the delayed response task. We were surprised by this finding, expecting before we initiated these experiments that we would need to affect much greater proportions of cortical neurons before behavioral impairments would become apparent. It is possible that the nature of neural coding in the spatial delayed response task, hypothesized to involve competition among microcircuits encoding possible goal locations (Arnsten et al., 2012) makes it particularly vulnerable to disruption of a small number of neurons in the network. This may not be the case for the involvement of other cortical/subcortical structures, in terms of the extent of disruption that would be required to impair behavior. For example, spatial navigation functions of the hippocampus apparently can be supported with only a small “minislab” of intact hippocampal tissue (Moser et al., 1995), suggesting that a much greater degree of transduction within the hippocampus would be required to produce behavioral deficits via an inhibitory DREADD receptor mechanism. Purely as an experimental design consideration, it will be critical, as in other lesion/inactivation studies of behavior, to take advantage of control tasks whose sensitivity to target structures is known, as well as dissociation methodology (Olton, 1991) to guard against errors in interpretation of functional deficits following neuronal inactivation (or activation). It is critical moving forward with these chemogenetic techniques that we proceed with both optimism and caution and support future findings with rigorous means of quantification to better verify these methods. As these methods develop, it will be important to continue to evaluate them in parallel with other approaches to interfering with neuronal activity, including pharmacological inactivations and permanent lesions, in order to obtain convergent evidence on the functions of neural systems for complex behavior in the primate brain.

## Acknowledgments

We are grateful to Scott Russo, Eric Nestler, and Bryan Roth for advice, to Jian Jin for synthesizing CNO HCl and Compound 21, and the NIMH Chemical Synthesis and Drug Supply Program for providing CNO. This work was supported by the Friedman Brain Institute at the Icahn School of Medicine at Mount Sinai as well as by NIH grant R21-NS096936.

**Author Contributions**
Designed Research: PRH, PGFB, PLC, PHR, MGB
Performed Research: NAU, SWB, WS, CGD, PGFB, PLC, PHR, MGB
Analyzed Data: NAU, SWB, WS, CGD, MGB
Wrote Paper: NAU, SWB, WS, PLC, PHR, MGB
The authors declare no competing financial interests.

